# Enabling Genotyping of Low-Biomass Zebra Mussel Samples via Multi-Displacement Amplification (MDA)

**DOI:** 10.64898/2026.11.16.767020

**Authors:** Víctor H. Hernández Elizárraga, Lindsey G. O’Brien, Michael A. McCartney, Daryl M. Gohl

## Abstract

Next-Generation Sequencing (NGS)-based genotyping enables population genomic studies of the history of biological invasions, allowing for inference of source populations and routes of spread. Methods for genotyping include whole genome sequencing (WGS), as well as lower-cost targeted approaches, such as microarrays, hybrid capture, or Genotyping by Sequencing (GBS). Using GBS, we previously genotyped ∼2,000 adult zebra mussels collected from over 100 populations across Minnesota, the Great Lakes, and the Upper Mississippi River. The need for sufficient template DNA restricts such genotyping approaches to adult animals, while genotyping lower biomass samples, such as zebra mussel veliger larvae remains challenging. Here, we applied Multiple Displacement Amplification (MDA) to increase DNA concentration from low-biomass samples. We demonstrate that MDA yields reproducible WGS and GBS data from as little as 1 ng of starting material and genotyped single veliger larvae from MDA amplified DNA, accurately assigning them to their known source populations. Furthermore, we showed that inheritance patterns can be tracked in MDA amplified DNA from veliger larvae and demonstrated an approach for forensic genetics of juvenile zebra mussels that were found contaminating “moss balls” and sold in the aquarium pet trade. Pairing MDA with high-resolution genomic workflows overcomes biomass limitations and represents a powerful toolkit to enhance early detection, trace invasion pathways, and mitigate the spread of the invasive zebra mussel. This approach is readily generalizable to other organisms or developmental stages where limiting quantities of DNA are available.

## Introduction

The invasive dreissenid mussels, the quagga mussel *Dreissena bugensis* (Adrusov 1897) and the zebra mussel *Dreissena polymorpha* (Pallas 1771), are listed as injurious species by the U.S. Fish and Wildlife Service and recognized as a biological threat by the U.S. Geological Survey^1,2^. In 2022, the estimated total cumulative cost of invasion, associated with mitigation costs and loss of revenue for quagga and zebra mussels, reached $46.4 billion and $19.3 billion, respectively^3^. These costs continue to grow. Although research aimed at designing effective control tools for dreissenid mussels is ongoing^4^, open-water control options are few. Understanding the life history and biology of these biofoulers, and improving methods for early detection are critical for developing effective control strategies.

The zebra mussel is a freshwater bivalve mollusk characterized by a triangular shell marked with alternating dark and light bands. This fecund spawner initiates its life cycle when water temperatures rise in early season, prompting the release of eggs and sperm into the water column, where fertilization occurs^5^. After this, the zygotes develop into typical bivalve planktonic larvae known as “veligers,” which possess a ciliated velum used for locomotion and feeding^6^. This free-swimming phase lasts for several weeks and facilitates dispersal by water currents^7^. Upon reaching the pediveliger stage the larvae develop a foot that searches and detects cues for suitable hard substrates for settlement (e.g rocks,vegetation, adult zebra mussels). Next, the settling larvae metamorphoses into a juvenile mussels secreting proteinaceous byssal threads to firmly attach itself. Juveniles then rapidly mature into filter-feeding adults, typically reaching sexual maturity within as little as a year^8,9^.

Native to the Ponto-Caspian region of Eurasia, zebra mussels were inadvertently introduced to the Great Lakes in the mid 1980s, likely via the discharge of ballast water from cargo ships^10^. Their populations spread rapidly due in large part to high fecundity, planktonic veliger larval dispersal, and byssal attachment leading to dense encrustations of natural and manmade substrates. Within a decade, zebra mussels expanded throughout all five Great Lakes and into major rivers, like the Mississippi and Hudson^11^. To date, the zebra mussel invasion costs the U.S. over $1 billion dollars per year in mitigation costs and lost revenue^3,12^. These mussels attach to water intake pipes in power stations, nuclear reactors, and drinking water utilities, and encrust dams, bridges, and other infrastructure. In addition, the presence of zebra mussels also impacts recreation, water-based transport, and property values through attachments to boats, docks, and shorelines, causing substantial costs in cleaning and removal. Concurrently, the ecological impacts include altering food webs by shunting carbon from pelagic to benthic pathways^13^ and decreasing the abundance and diversity of other planktonic and macrobenthic species^14^, harming ecosystems by smothering some native species and by outcompeting others for food and space^15^.

Genomic surveillance has revolutionized the detection and study of invasive species by applying highly sensitive molecular detection tools, such as DNA sequencing, DNA microarrays^16,17^, and the analysis of environmental DNA (eDNA)^18,19^. The latter involves targeted, high-sensitivity detection of specific genomic sequences in the environment using quantitative PCR or high-throughput DNA (and recently, RNA) sequencing to screen entire communities simultaneously for high-impact exotic species. Additional sequence-based genotyping methods analyze genomic variants to trace invasion pathways and identify source populations, and to study how invasive species adapt genetically to their new environments.

Genotyping-By-Sequencing (GBS) is an approach that enables the simultaneous discovery of thousands of DNA markers; primarily single nucleotide polymorphisms (SNPs)^20^. This approach is part of a suite of methods that rely on genome complexity reduction^21^, achieved by digesting genomic DNA (gDNA) with specific restriction enzymes to fragment the genome at conserved cut sites (Figure 1A). Adapters are ligated to these fragments, ligation products are amplified with sample-specific indexing primers, allowing samples from hundreds of individuals to be multiplexed and sequenced together in a single sequencing run. To identify SNPs, the resulting sequence reads may be mapped (aligned) back to a reference genome, or inferred *de novo*. This reduced-representation sequencing approach lowers the cost significantly compared to WGS (Figure 1B) while providing tunable marker density for specific applications. GBS is particularly effective for studying non-model organisms and species with large, complex genomes.

**Figure 1.**
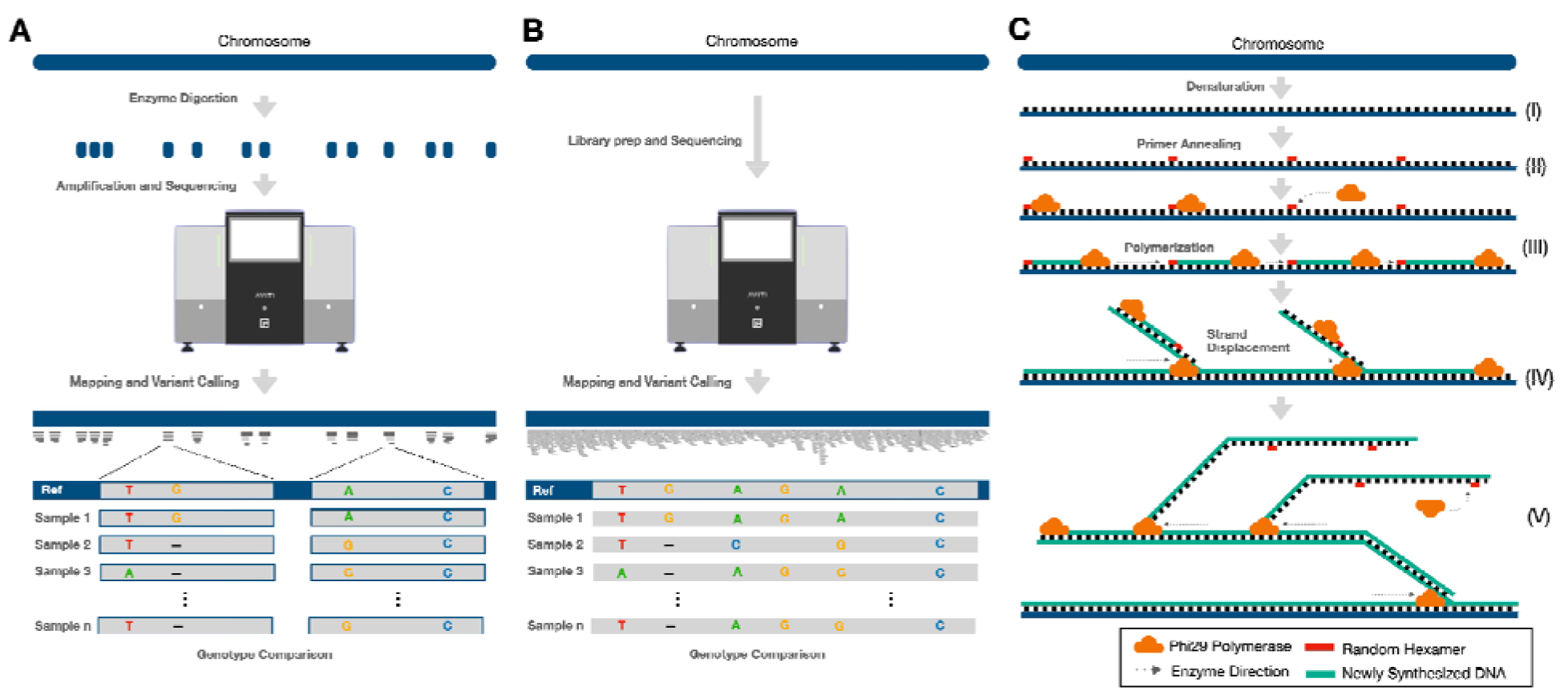
Overview of the Genotyping workflow and Multi Displacement Amplification (MDA) reaction mechanism. A) GBS analysis begins with enzyme digestion to reduce genome complexity. Here, gDNA is fragmented at conserved restriction sites. Following adapter ligation and barcoding, these fragments undergo amplification and sequencing. The resulting reads may be mapped to a reference genome (or called *de novo*) and subjected to variant calling. The final genotype comparison identifies allelic variations, including Single Nucleotide Polymorphism (SNPs). B) The whole genome sequencing (WGS) genotyping workflow begins with preparing libraries from mechanically or enzymatically sheared genomic DNA (gDNA), followed by end-repair of fragments and adapter ligation for sequencing. Tagmentation libraries for WGS combine these into one step, in which a transposase cuts gDNA and tags fragments in one reaction. Unlike GBS, WGS is not a “reduced representation” approach. WGS sequencing reads can survey the entire genome or targeted subsets (e.g. the exome), whereas GBS surveys variant only in regions adjacent to restriction sites. C) Representation of the Multiple Displacement Amplification (MDA) Reaction. (I) Physical denaturation of the gDNA template. (II) Annealing of random hexamer primers to the denatured strands. (III) Initiation of DNA synthesis by the high-fidelity Phi29 polymerase. (IV) Strand displacement synthesis, where the polymerase displaces the preceding strand while extending the new one. (V) Exponential amplification through secondary priming on displaced strands, resulting in high-molecular-weight branching structures and a substantial increase in DNA yield.

We previously genotyped ∼2,000 adult zebra mussels collected from over 100 populations across Minnesota, the Great Lakes, and the Upper Mississippi River using GBS. This study identified over 5,500 SNPs, which allowed us to identify Lake Superior, the Upper Mississippi, and inland “hub” lakes as sources of zebra mussel infestation^22^. A major goal of this and our ongoing work is to provide natural resource managers with a tool to rapidly identify lakes that are responsible for the spread of zebra mussels. In the few cases where data from intensive annual and seasonal sampling are available, early-stage invasions are first identified through the detection of zebra mussel veliger larvae in plankton tows^26^, and prior to the discovery of adult mussels. Of the nine largest walleye fishing lakes in the Minnesota DNR’s Large Lakes Program five are now zebra mussel infested. Veligers were detected first in four of these lakes, and in the fifth (Mille Lacs Lake) veliger sampling was not done until after adults were discovered^23,24^. An additional case is Lake Simcoe in Ontario, Canada. Here, intensive study of early life stages (veligers in the plankton, post-veligers and juveniles newly settled onto artificial substrates, juvenile and adult mussels collected from natural substrates) confirmed the first appearance of larvae, post-larvae, and mussels in the same year (1994)^25^. Genotyping such samples is difficult, as veligers typically yield around 1 nanogram (ng) of DNA or less. WGS and GBS analysis typically requires on the order of tens to hundreds of ng of gDNA, which precludes their use on single-veliger samples when processed using current methods.

MDA (Figure 1C), a whole-genome amplification technique, increases small amounts of DNA in biological samples to enable downstream analyses^27^. The experimental MDA workflow begins by heating to physically denature the DNA. Subsequently, a gradual decrease in temperature allows random, exonuclease-resistant primers to stochastically bind to regions across the entire genome. The addition of the high-fidelity and highly processive Phi29 polymerase, along with Bovine Serum Albumin (BSA) as a stabilizer, facilitates the isothermal amplification (at 30°C) of ∼12 kb genomic amplicons^28^. During the reaction, multiple strand displacement events occur; the polymerase first copies the gDNA template and then synthesizes copies of the newly formed strands while displacing the previously extended ones^29^. MDA can generate micrograms of DNA from as little as a few femtograms (10^-6^ ng), providing sufficient material for Next-Generation Sequencing (NGS) technologies.

MDA has been applied to a wide array of research areas, including single-cell sequencing^28,30^, human genome amplification^31^, molecular epidemiology^32^, barcoding amplification in spatial genomics^33^, amplification of complex DNA mixtures^34^, and metagenomic analyses^35^. In this study, we evaluated the integration of MDA with GBS and WGS workflows to overcome the gDNA input limitations associated with low-biomass samples, including single-veliger larvae, and demonstrated the versatility of this approach under different scenarios. We aim to provide a reliable genomic method for detecting and tracking early-stage zebra mussel invasions, ultimately delivering the time-sensitive data required to inform management strategies aimed at curbing the spread of this damaging invasive species.

## Methods

### Adult Zebra Mussel and Plankton Tow Collections

Zebra mussels were collected by SCUBA and snorkeling at Gray’s Bay (44.9509444°, - 93.49855556°), Lake Minnetonka, Minnesota (MN), USA, from bridge pylons and riprap along the lake bottom (Prohibited Invasive Species Permit #826, valid from 10/24/2023 through 12/31/2028; MN of Natural Resources (DNR) Division of Ecological and Water Resources). Two mesh collection bags were used to target different size classes: (1) a white fine-mesh laundry bag with 0.5-mm mesh openings for mussels <10 mm, and (2) a black standard sports mesh bag with 5.0-mm mesh openings for mussels >2 cm. Mussels <10 mm were collected by retrieving riprap pieces and stones densely colonized by juvenile mussels from the lake bottom at approximately 4 m depth. Larger mussels (>2 cm) were collected directly from bridge pylons at depths ranging from 1–3 m. Mussels were carefully detached from the pylon surfaces using a plastic card, minimizing damage to the shells and byssal threads. Upon surfacing, divers placed mussels from each size class into separate insulated coolers containing ice, with a layer of damp newspaper placed over the ice to maintain moisture and reduce thermal stress during transport.

In addition, vertical plankton tows were conducted from a 14.5 ft boat at Gray’s Bay and Crystal Bay, Lake Minnetonka (44.953983°, -93.494867°). At each site, the deepest area of the bay was located, and the vessel was anchored prior to sampling. A simple plankton net (30 cm mouth diameter × 120 cm-long 50-µm mesh) equipped with a stainless steel ring and bridle and a two-piece PVC cod end with a detachable lower section containing a 50 µm mesh screen and ballast weight was used for collections. The net was deployed on a 12 m marked line with a reel and lowered vertically through the water column, taking care to avoid contact with the lake bottom, then retrieved at a controlled rate. Following retrieval, the net was rinsed externally with lake water to ensure all collected material was flushed into the cod end. Cod end contents were transferred into a 250 mL Nalgene bottle prefilled with 175 mL of 100% ethanol, and the cod end screen was rinsed with lake water to bring the total volume to 250 mL.

### DNA from Fresh and Banked Mussel Samples

gDNA was extracted from adult mussel tissue collected from Lake Minnetonka using the DNeasy Blood and Tissue Kit (QIAGEN, Germantown, MD) according to the manufacturer’s protocol. Banked gDNA was also utilized from Duluth-Superior Harbor (MN), the Illinois River (IL), and Lake Erie (MI) adult specimens originating from our previous genotyping study^22^. For larval samples, zebra mussel veligers were isolated from plankton tows and identified using cross-polarized microscopy^36^. This method utilizes the birefringence of the crystalline calcite structure in veliger shells, which produces a distinct “X” diffraction pattern under cross-polarized light. Following visual confirmation, gDNA was extracted from single veligers using the DNeasy kit and protocol described above. The concentration of the isolated and banked gDNA was verified using the Qubit High Sensitivity dsDNA Assay Kit (Thermo Fisher Scientific, Waltham, MA).

### Extraction of Genomic DNA from Drosophila Stocks

In addition to adult mussel and veliger samples, gDNA was extracted from several *Drosophila melanogaster* laboratory stocks, specifically IsoD1 (an isogenized wild type strain)^37^, and a number of stocks obtained from the Bloomington Drosophila Stock Center: iso-1 (the Drosophila genome strain, RRID:BDSC_2057), OR-A (RRID:BDSC_99765), BK-T (RRID:BDSC_99749), F9-D (RRID:BDSC_99754), and CS-B (RRID:BDSC_99750). These extractions were performed using the same DNeasy Blood and Tissue Kit (QIAGEN, Germantown, MD) and protocol as described above.

### Standardization of MDA Parameters

MDA reactions were performed using Phi29 DNA polymerase [New England Biolabs, Ipswich, MA (NEB)] in a 20 µL final reaction volume. Initially, an annealing mixture, consisting of Exo-Resistant Random Primers (hexamers) (Thermo Fisher Scientific, Waltham, MA; 5 µM final), dNTPs (200 µM final), and a dilution series of template DNA (100, 10, 1, 0.1, 0.01, and 0.001 ng; n = 6), was incubated at 95°C for 5 minutes and then cooled to 4°C (cooling rate of 2°C/s). An amplification master mix containing Phi29 reaction buffer (NEB: 1X), BSA (100 µg/mL), and Phi29 polymerase was then added to the annealed template and incubated at 30°C for 15 hours. To evaluate amplification efficiency over time, reactions with 1 ng of input DNA were incubated at 30°C for 3, 6, 9, 12, or 15 hours.

All MDA products were treated with 1 µL of T7 Endonuclease I (NEB) in NEBuffer 2 and incubated at 37°C for 15 minutes to resolve branching structures (Figure 1C). DNA was purified post-reaction, via lithium chloride/isopropanol precipitation. In short, samples were diluted with 80 µL of molecular grade water, mixed with 120 µL of isopropanol and 20 µL of 3 M LiCl, and incubated on ice for 10 minutes. Then, the DNA was pelleted by centrifugation at 12,000 × g for 10 minutes, washed twice with 70% ethanol (centrifuging at 12,000 × g for 5 minutes), and resuspended in molecular grade water. Final yields were quantified using the Qubit High Sensitivity dsDNA Assay Kit.

To verify the integrity of the MDA amplified DNA, a PCR assay targeting the 28S rRNA gene was performed. Reactions were prepared in a 25 µL final volume using KAPA HiFi PCR reagents (Kapa Biosystems, Wilmington, MA.). Each reaction contained 1X KAPA HiFi GC Buffer, 0.3 mM dNTPs, 0.3 µM each of forward (5’-TCCGATAGCGCACAAGTACC-3’) and reverse (5’-TTGCACGTCAGAATCGCTAC-3’) primers, 0.5 µL of KAPA HiFi HotStart Polymerase, and 1 µL of template DNA. Thermal cycling was initiated with an initial denaturation at 94°C for 2 minutes, followed by 40 cycles of denaturation at 94°C for 1 minute, annealing at 46.5°C for 1 minute, and extension at 72°C for 1 minute 30 seconds, with a final extension step at 72°C for 5 minutes.

### GBS Analysis: DNA Digestion, Size selection, Sequencing and Variant Calling

Genotyping by Sequencing libraries were prepared and sequenced at the University of Minnesota Genomics Center. Briefly, gDNA was digested with *Sbf*I-HF and ligated to TGCA-overhang adaptors with T4 DNA ligase containing binding sites for indexing primers. Adapter-ligated DNA fragments were purified with solid phase reversible immobilization SPRI beads and amplified with NEBNext® High-Fidelity 2X PCR Master Mix with combinatorial dual indexing primers:

Forward indexing primer: AATGATACGGCGACCACCGAGATCTACACXXXXXXXXTCGTCGGCAGCGTC

Reverse indexing primer: CAAGCAGAAGACGGCATACGAGATXXXXXXXXGTCTCGTGGGCTCGG “X” represents sample-specific barcodes. Libraries were quantified with the Quant-IT PicoGreen dsDNA Kit (Thermo Fisher Scientific, Waltham, MA), pooled by mass, and purified using SPRI beads. The final library fragment size distribution was determined using a Tapestation High Sensitivity assay (Agilent, Santa Clara, CA). Libraries were sequenced on an Element AVITI sequencer (Element Biosciences, San Diego, CA) with a paired end (2x150 bp) kit.

Following sequencing, reads were demultiplexed, trimmed, and aligned to the zebra mussel reference genome^38^ (NCBI Accession No. GCF_020536995.1) using Bowtie2 v.2.5.4^39^. SNP markers were called using FreeBayes v.1.1.0^40^. The resulting variants were processed and filtered using PLINK v.1.90^41^. In brief, initial VCF files were converted to binary PED format while maintaining allele order (--keep-allele-order flag in PLINK). Variant IDs were assigned using standard format [--set-missing-var-ids @:# (where @ = chromosome code and # = SNP position in bp). To ensure high-quality genotypes, we filtered the dataset to retain only biallelic SNPs, applying a maximum per-variant missingness threshold of 20% (--geno 0.20) and a maximum per-sample missingness threshold of 50% (--mind 0.5). Finally, to minimize the effects of linkage disequilibrium, SNP pruning was performed using an independent pairwise search with a 200-variant window, a step size of 1 SNP, and a threshold value for the squared genotypic correlation between pairs of SNPs (r2, without phasing) = 0.5.

### Population Structure Calculations and Principal Component Analysis (PCA)

Genotypic data filtering and linkage disequilibrium (LD) pruning were conducted using PLINK (v1.9)^41,42^ and options set to--snps-only, with a minor allele frequency ≥0.01 (--maf 0.01), a SNP missingness threshold of 5% (--geno 0.05), and genotype missingness threshold for individuals of 50% (--mind 0.5). SNPs in linkage disequilibrium were pruned were pruned using the sliding window settings described earlier (--indep-pairwise 200 1 0.5) Population structure and ancestry estimation were evaluated using ADMIXTURE^43^ on the LD-pruned SNP set. Principal Component Analysis (PCA) was conducted using PLINK (v1.9)^41,42^. Eigenvectors and eigenvalues were calculated, using the -pca flag, to evaluate population structure and sample relationships across the primary principal components. Genetic distances were calculated from the filtered VCF files using the bitwise.dist function in the R package poppr^44^ (v2.9.) via a converted genlight object. A neighbor-joining phylogenetic tree was generated using the BioNJ algorithm (bionj) implemented in ape^45^. Node support was assessed by performing 1,000 bootstrap replicates with the aboot function. The resulting consensus phylogenetic tree was saved in NEXUS format and visualized using iTOL^46^ software v7. Genomic distances were calculated in PLINK v.1.9^41^ using the Identity-by-State (IBS) method. First, the VCF was recoded into a pedigree format while allowing for extra-chromosomal scaffolds (--allow-extra-chr). Subsequently, a distance matrix was generated using the --cluster and --matrix flags to quantify the genetic similarity between individuals. The resulting IBS matrix was then used to construct a dendrogram for visualization using a custom script in R.

### MDA Experimental Applications

Following the MDA optimization process described above, an input concentration of 1 ng (except when indicated otherwise) and a reaction time of 12 hours were selected for the following experimental designs.

### Effect of MDA on WGS Output

To determine the impact of MDA on sequencing performance, gDNA from *Drosophila melanogaster* (n = 6) and adult zebra mussel (n = 6) samples were diluted to 1 ng and amplified via MDA. The resulting MDA products, along with their corresponding undiluted genomic controls, were sequenced at the UMGC. Twist FlexPrep was used to prepare Drosophila libraries and zebra mussel libraries were prepared with Ultima PCR-Free library prep and both sets of samples were sequenced on the Ultima Genomics UG100 sequencer. For Drosophila samples, whole-genome reads were aligned to the reference genome (NCBI Accession No. GCA_000001215.4) using the Ultima Aligner (ua) v.2.2.1^47^. Variant calling was performed using Ultima make_examples v.3.1.10^47^ and Ultima call_variants v.2.2.4^47^. The quality of the sequencing and alignment data was assessed using Picard v.2.25.6^48^ via the CollectWgsMetrics and CollectMultipleMetrics tools, respectively. To identify integration elements, sequences were matched to reads using SeqKit v.2.9.0^49^, and subsequent alignments to the reference genome were performed with Bowtie2 v.2.5.4^39^ . The resulting output files in SAM format were converted to BAM format using SAMtools v.1.20^50^ and then to BED format using BEDtools v.2.31.1^51^. Finally, filtered BED files were generated by excluding sites with a mapping quality (MAPQ) score of less than 20 to ensure high-confidence genomic localization. For zebra mussels, demultiplexing of reads, trimming adapter sequences, and alignment to the reference genome was performed on the UG100. Variant calling of the CRAM alignment files was completed using the Ultima Genomics-adapted DeepVariant variant calling software (make_examples version 3.1.10; call_variants version 2.2.4; https://hub.docker.com/u/ultimagenomics). Variant call file statistics were tabulated using bcftools version 1.21^50^ on the vcf files subset to variants with quality scores of q20 or higher. Counts of variant types were tabulated from the vcf files subset to variants with quality scores of q20 or higher.

### Effect of MDA on Population Genotyping

To assess the effect of MDA on the genotyping accuracy of adult mussels from diverse locations that span the range of among-population variation we have found to date^22^, gDNA extracts from Duluth/Superior Harbor on Lake Superior, the Illinois River, and Lake Erie (n = 3 individuals per location, each individual with three technical replicates) were diluted to 1 ng and amplified using the MDA conditions described above. Both amplified and unamplified materials (54 samples total) were analyzed using GBS at the UMGC with the method described above.

### Single-Veliger Genotyping

To evaluate the feasibility of generating high-resolution genotyping data from a single larva, ∼1 ng of gDNA from Lake Minnetonka veligers (n = 6) was amplified using MDA. These samples were analyzed via GBS alongside unamplified gDNA from 10 adult mussels from the same location to serve as a reference. Typically, template DNA is not amplified prior to digestion and ligation steps in the GBS workflow.

### Genetic Trio Analysis

One male and one female zebra mussel was identified by gamete release through serotonin-induced spawning^52^. Mussels (>10 mm) were first scrubbed with 10% bleach, neutralized with sodium thiosulfate, and depurated in artificial fresh water (AFW) for one hour. Spawning was induced by immersing each single mussel in a 10 mM serotonin creatinine sulfate solution via individual induction in a 50 mL centrifuge tube. After induction and gamete release, males were kept refrigerated (to prevent further spawning), while females were transferred to fresh artificial fresh water (AFW) to complete oocyte release. Eggs were cleaned of debris through serial sedimentation and washing, then concentrated into a 2% v/v suspension in AFW. Sperm was collected from fragments of testis macerated in AFW and checked for live sperm using a “squash” under a cover slip on a microscope slide observed under 200X. Fertilization was achieved by inoculating the egg suspension with a calculated volume of sperm to achieve a target ratio of approximately 50 oocytes per 200,000 sperm cells. Successful fertilization was verified by monitoring polar body formation and two-cell stage cleavage^53^ before transferring embryos to room temperature AFW in glass jars. DNA was extracted from the two parents and six young veliger larvae offspring (∼5 days old) using the DNeasy Blood and Tissue Kit (QIAGEN, Germantown, MD). Larval DNA (∼0.01 ng; n=6) was subjected to MDA, while parental DNA (n=2) was processed without MDA. These samples were subsequently analyzed using GBS.

### Case Study: Moss Ball Mussels

In 2021, zebra mussels were found attached to commercial aquarium moss balls (a rare form of the green alga *Aegagropila linnaei* sold in the pet trade. The U.S. Geological Survey (USGS) worked alongside the U.S. Fish and Wildlife Service (USFWS) and pet stores to remove the contaminated moss balls from stores^54,55^. Four DNA samples, derived from PCR-confirmed zebra mussel attached to moss balls, were shipped to our laboratory for genotyping by the USGS Upper Midwest Environmental Sciences Center. In parallel, eight moss ball samples were sent to the Minnesota Aquatic Invasive Species Research Center (MAISRC) at the University of Minnesota. Upon inspection of these samples, we found small mussels (∼2 mm) and shells attached to the moss balls. DNA was extracted from these samples using the aforementioned method. Due to low DNA yield, all samples were subjected to MDA whole-genome amplification and genotyped using GBS.

## Results

### MDA Reactions Yield High-Concentration DNA Products

MDA of genomic DNA extracted from zebra mussels generated high molecular weight template DNA (Figure 2A). PCR amplification of the 28S gene from this MDA-treated DNA (Figure 2B) produced high product yields. Within the range of input MDA-DNA assessed (0.001–100 ng), a plateau effect was observed at approximately 2500–3000 ng of output mass, representing the physical saturation limit of the reaction (Figure 2C). Our results showing that the steepest part of the curve falls between between -3 and -1 Log(ng) (1 to 100 pg), indicates that the protocol is highly effective at amplifying very small amounts of DNA. This indicates that the method is suitable for zebra mussel larval samples, as we found the DNA yield from each veliger larva to average about 1 ng (x = 1.1 ± *s* = 0.065 ng). To determine the optimal reaction time, we evaluated a window of 3–15 hours at three-hour intervals using 1 ng of input material and found 12 hours to be sufficient to reach the plateau, while a minimum of 9 hours is necessary to observe the exponential amplification phase (Figure 2D). Based on these results, we established optimal reaction conditions for whole genome amplification of mussel adults and larvae at an input mass of 1 ng and a 12-hour reaction time.

**Figure 2.**
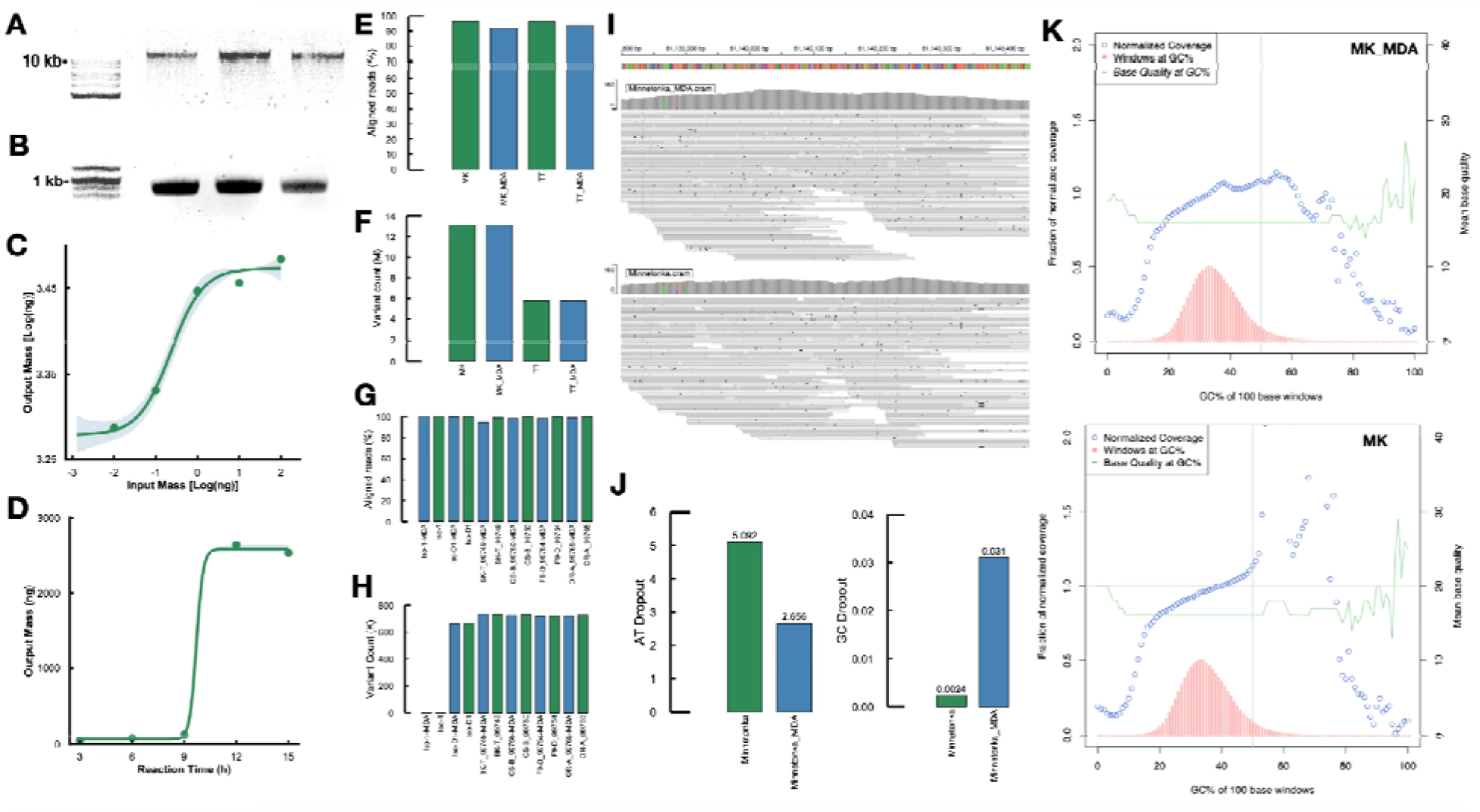
Multi-displacement Amplification (MDA) Optimization and whole-genome sequencing (WGS) validation of the MDA protocol. (A) Representative gel electrophoresis image showing MDA-amplified high-molecular-weight gDNA from zebra mussels. (B) PCR amplification products of the zebra mussel 28S rRNA gene using MDA-treated DNA as a template. (C) MDA output mass across input DNA masses. (D) MDA output mass across reaction incubation times. (E, F) Comparison of WGS metrics between control and MDA-treated adult zebra mussel samples from Minnetonka (MK) and Turtle Lake (TT), showing (E) percentage of aligned reads and (F) total variant counts (in millions). (G, H) Validation using six Drosophila reference stocks with and without MDA treatment, showing (G) percentage of aligned reads and (H) total number of variants (blue = MDA, green = unamplified). (I) Representative section of the zebra mussel genome (Minnetonka samples) showing mapped reads for both MDA-amplified and untreated DNA. (J) AT and GC dropout percentages for the Minnetonka population. (K) Sequencing GC bias and base quality across 100 bp genomic window for Minnetonka samples. Red bars indicate genome GC content. Blue circles represent the fraction of normalized coverage. Green line shows base quality at GC.

### Effect of MDA on WGS from Diluted Zebra Mussel Adult DNA

To assess the uniformity of whole genome amplification and to examine the amount of contaminating DNA being amplified by MDA, we performed WGS on both MDA-treated and control samples from adult zebra mussels and Drosophila. Raw reads generated from the zebra mussel samples from Minnetonka (MK) and Turtle Lake (TT) displayed similarly high-quality (Q-scores >30) across both control and MDA-treated groups. We observed an approximate 3% decrease in the percentage of mapped reads in the MDA-treated samples relative to unamplified samples (Figure 2E). Nevertheless, these samples still showed >93% mapping, indicating that the vast majority of the sequenced DNA originated from zebra mussels. We determined the total number of variants, including single- and multi-nucleotide polymorphisms, and identified no notable differences between the control and treated DNA samples (Figure 2F). In parallel, six DNA samples from Drosophila strains were treated with MDA, and the same genome sequencing metrics were evaluated. In this reference material, MDA did not modify the alignment rates (Figure 2G), or total variant counts (Figure 2H). The iso-1 strain corresponds to the stock used to generate the Drosophila reference genome and as expected this had very few variants relative to the reference genome. A representative section of the zebra mussel genome showing comparable coverage of mapped reads for both MDA-amplified and untreated DNA is depicted in Figure 2I, while the overall of AT and GC dropouts, which indicate the percentage of misaligned reads linked to regions with low GC content (less than 50%) or high GC content (more than 50%), respectively. for the Minnetonka population are shown in Figure 2J. Specifically, the Minnetonka-MDA sample displayed similar sequencing performance in the middle region corresponding to 20% to 50% GC (Figure 2K).

### Genotyping Data Quality in MDA-treated Adult Zebra Mussel DNA

To examine the effect of MDA on the genotypes of three zebra mussel populations (Duluth Harbor, Illinois River, and Lake Erie), high-quality gDNA was diluted to 1 ng and treated with MDA (Figure 3A). Slight differences in guanine-cytosine (GC) content and the percentage of reads remaining after deduplication were observed between control and MDA-treated samples. However, mapping the generated datasets against the zebra mussel reference genome resulted in consistent alignment rates (>95%) between groups (Figure 3B). Post-mapping variant calling showed that MDA-treated samples exhibited low fractions of missing genotype data and slightly lower total number of variants, including all types of single- and multi-nucleotide variants. It is worth noting that one replicate (LE2_M_2) exhibited a drop in the percentage of aligned reads; however, the remaining two technical replicates were unaffected and displayed adequate alignment rates, likely indicating a failed library preparation for this sample. The MDA samples remained consistent in the total number of SNPs per sample (Figure 3C). To determin sample clustering patterns, the genetic distance between samples, calculated using the BioNJ method (Figure 3D), IBS method (Supplementary Figure 1), and the PCA analysis (Figure 3E) using control and treated mussel populations displayed compact clusters with their respective MDA-treated samples, indicating high reproducibility between technical and biological replicates.

**Figure 3.**
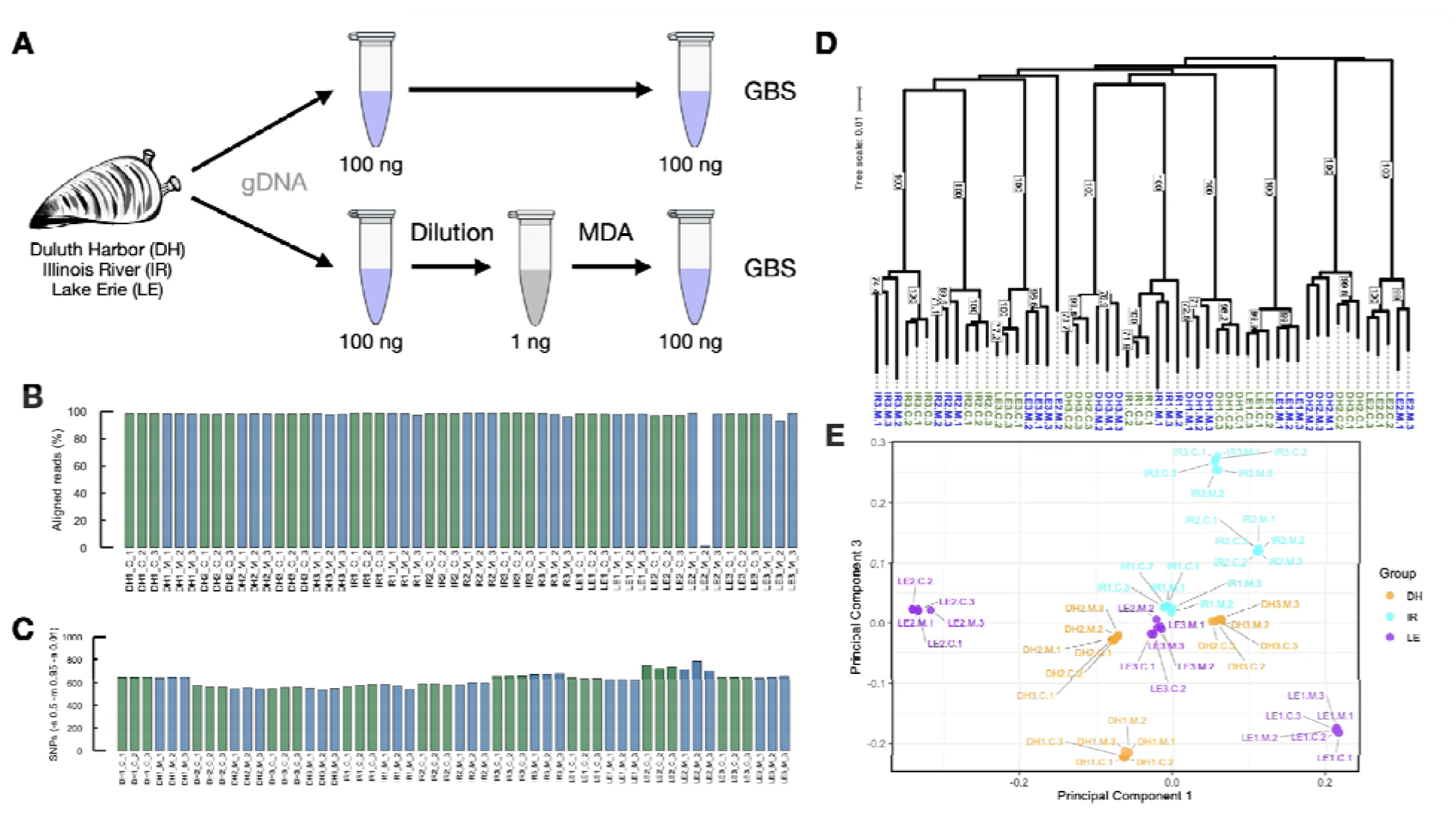
MDA performance on adult zebra mussel gDNA quality and genotypic data. (A) Experimental workflow for comparing control gDNA (100 ng) and MDA-treated samples (1 ng input) across three populations (Duluth Harbor, Illinois River, and Lake Erie) using Genotyping-by-Sequencing (GBS). (B) Alignment rates against the zebra mussel reference genome. (C) Total SNP counts per sample, indicating high concordance in variant detection between treatment (blue = MDA; green = unamplified). (D) Biological Neighbor-Joining (BioNJ) phylogenetic tree generated based on genetic bitwise distances. Circles at nodes indicate bootstrap support value based on 1,000 replicates; control (C, green) and MDA-treated (M, blue) replicates from the same biological source. (E) Principal component analysis (PC1 vs. PC3). Sample populations are indicated by color: DH = Duluth Harbor (orange), IR = Illinois River (cyan), and LE = Lake Erie (purple). Sample labels indicate processing condition (C = unamplified control; M = multiple displacement amplification [MDA]).

### MDA Enables Generation of Genotypic Data from Single Veliger Larvae

To evaluate the feasibility of generating genotyping data from larvae, we analyzed MDA-treated genomic material from Minnetonka veliger samples using GBS and compared the output with adult mussels from the same location (Figure 4A). Similar to our observations for amplified adult DNA, sequencing quality scores remained high for the larval MDA-DNA. Comparable GC content and deduplication percentages were observed between the adult and veliger datasets. The high percentage of mapped reads (>95%) (Figure 4B) confirms that this protocol successfully amplifies zebra mussel genomic sequences that align well to the reference genome, ruling out the possibility of significant amplification of non-target or contaminant DNA. Additionally, variable fractions of missing genotypes and total variant counts were observed between adult and larval samples. Consistent with results from zebra mussel adults, a less variable number of SNP variants were detected in the MDA-treated larval samples (Figure 4C). Ultimately, we utilized pruned genotype data, consisting exclusively of SNPs, to compare the genomic distances between the three control populations (Duluth Harbor, Illinois River, and Lake Erie) and the Minnetonka adult and larval samples using the BioNJ algorithm and the IBS method. The dendrograms in Figure 4D and Supplementary Figure 2 illustrate the sample clustering for this comparison. As expected, the control populations formed distinct, population-specific clusters. Notably, most of the Minnetonka adults (MK) and veligers (MK_V) grouped together within a single cluster as depicted in Figure 4D, with the exception of a few samples whose branches had low bootstrap support (<70%). This shows that genotyping data generated from MDA-treated zebra mussel larval samples can be used to identify population-specific SNP signatures and trace relatedness across the tested populations.

**Figure 4.**
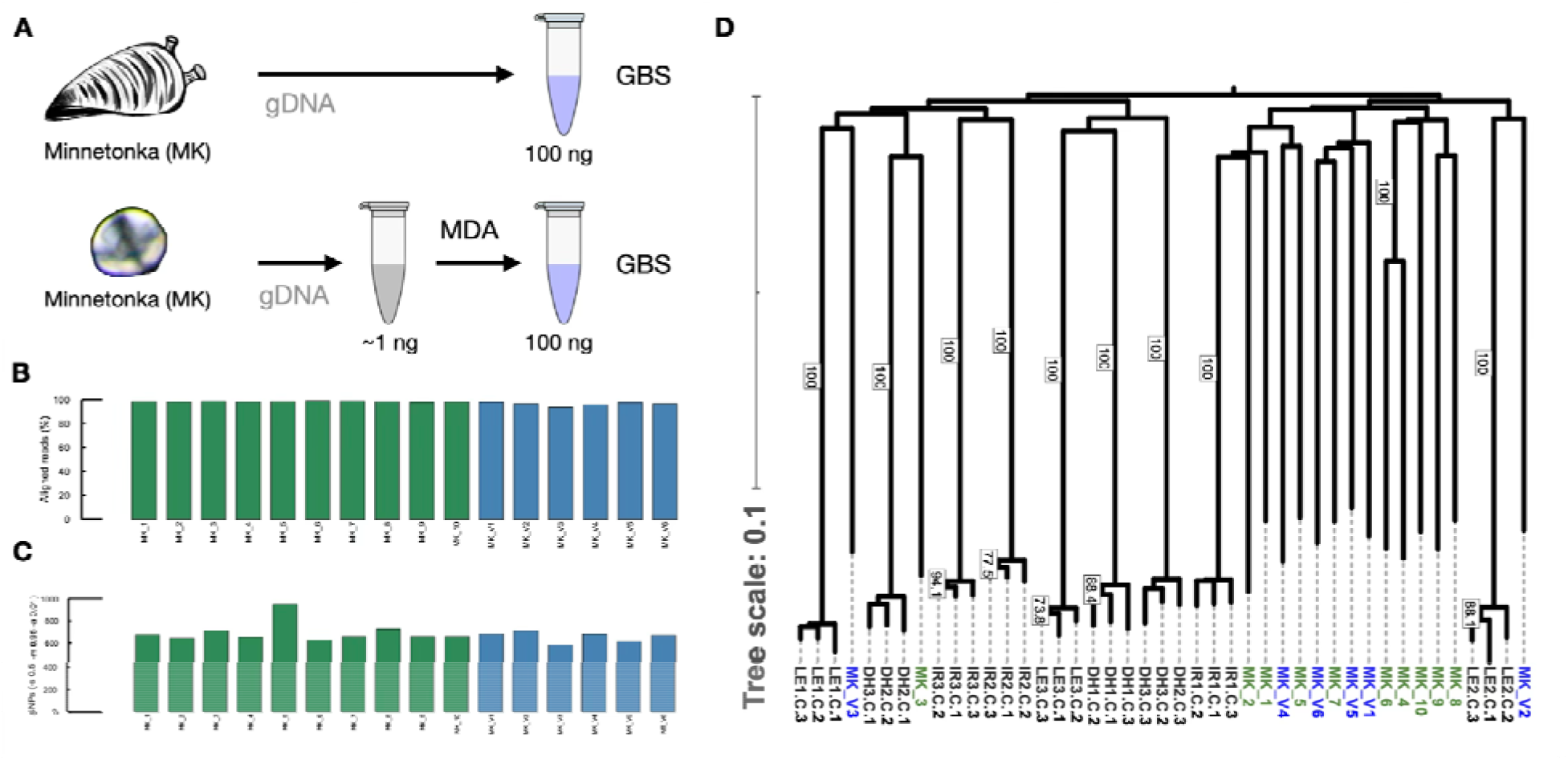
Comparison of MDA-GBS output and genetic clustering between adult and singl veliger larvae. (A) Experimental design comparing control gDNA from Minnetonka adults to MDA-amplified gDNA from single Minnetonka veliger larvae. (B) Alignment rates to the zebra mussel reference genome. (C) SNP counts across adult and larval samples (blue = MDA larvae, green = unamplified adults, grey = outgroup adults). (D) BioNJ phylogenetic tree generated based on genetic bitwise distances. Circles at nodes indicate bootstrap support values based on 1,000 replicates; the dendrogram shows Minnetonka (MK) adults and veligers (MK_V) alongside the three reference populations from Figure 2.

### MDA Allows Detection of Inheritance Patterns

To further confirm that MDA derived genotypes accurately reflect the genetic makeup of zebra mussel larvae, we carried out a “trio” sequencing experiment where expected larval SNPs could be inferred from parental genetic makeup. We analyzed MDA-treated DNA from veliger larvae in comparison with parental genotypes (Figure 5A). Figures 5B through 5E depict the successful spawning and *in vitro* fertilization processes. Light microscopy images of viable zebra mussel sperm, and eggs are shown in Figures 5B and 5C, respectively. Fertilization was confirmed by polar body formation and cleavage at the two-cell and multiple-cell stages, as displayed in Figure 5D (see arrow). Finally, a representative image of a young veliger larva offspring (∼5 days old) is shown in Figure 5E. GBS analysis of zebra mussel offspring revealed low alignment rates, with high missing genotype fractions for some samples. A higher number of variants wa observed, particularly for samples Offspring_V1 and Offspring_V3 (Supplementary Figure 4). Due to current limitations with culturing veligers in the lab, these larvae were from earlier stages and smaller than those genotyped in Figure 4. These samples yielded a lower DNA concentration (∼0.01 ng) than wild-caught veliger samples, which likely resulted in the more variable sequencing performance. Overall, SNP counts ranged from 400 to 600 for both parents and offspring (Figure 5F); this range is consistent with the number of SNPs detected in adult sample from the DH, IR, and LE populations both with and without MDA (Figure 3C), as well as in adults and veliger larvae from Lake Minnetonka (Figure 4D). Genetic inheritance patterns were identified across several genomic regions. Figure 5G displays the chromosomal region Chr4:121,160,675–121,160,817. In the zebra mussel reference genome, this corresponds to a site where SNPs were detected; nucleotide letters represent the reference sequence, and the piled sample names show the genotype at each position. For this particular example, the male and female were identified as homozygous (gray) or heterozygous (blue) relative to the reference. As expected, sites with homozygous parental alleles produced homozygous offspring, while sites with heterozygous parental alleles produced either homozygous or heterozygous offspring consistent with expected patterns of Mendelian inheritance.

**Figure 5.**
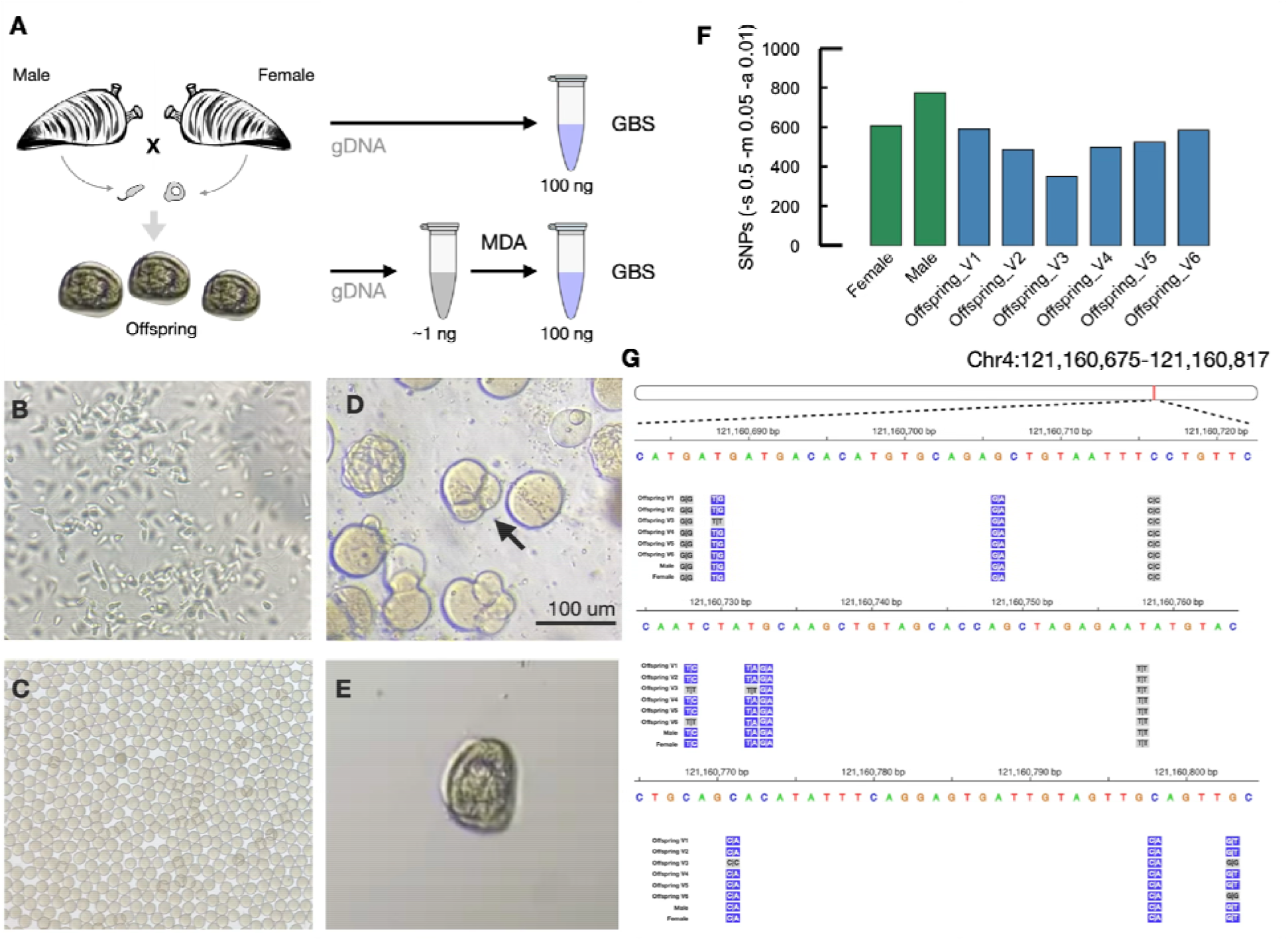
MDA-enabled detection of inheritance patterns in zebra mussels. (A) Experimental workflow showing the cross between a male and female zebra mussel to produce veliger larvae offspring. (B–E) Representative light microscopy images of the developmental stages: (B) viable sperm, (C) eggs, (D) fertilization confirmed by cell cleavage, and (E) a 5-day-old veliger larva. (F) Final SNP counts after filtering. (G) Visualization of a ∼130 bp chromosomal region on Chr4 (121,160,675–121,160,817). Nucleotide letters represent the reference sequence, and blue boxes indicate heterozygous calls and gray boxes indicate homozygous calls relative to the reference.

### Zebra Mussels Found in Pet Store Aquarium Moss Balls

MDA was used to generate genotyping data from zebra mussels found in imported moss balls from pet stores (Figure 6A). We analyzed DNA from samples obtained both by the USGS and MAISRC. Overall, these samples showed a low alignment rate and a high fraction of missing genotypes. The samples from MAISRC were originally confiscated in 2021, frozen in water, and processed for this study four years later. The DNA received from the USGS was extracted from archived mussels. Prior to DNA extraction, these samples were subjected to two freeze-thaw cycles: specimens were initially frozen within damp moss balls, thawed to separate the mussels from the substrate, and subsequently refrozen without liquid media or preservatives. Storage in water for months or years, together with freeze-thaw cycles, could compromise the integrity of the samples through cell lysis and loss of mussel DNA or through cross-contamination with DNA from other organisms in the moss ball samples. This may explain the low mapping rate observed in the resulting GBS data. Nevertheless, from these 12 moss ball zebra mussel samples, we identified 1,207 markers across 248 loci. Genomic distance analysis of these samples, using DH, IR, and LE as outgroup references, showed that the USGS samples (Figure 6B and Supplementary Figure 3), had a lower missing genotype fraction, did not cluster with the control populations and formed a separate clade. The samples from MAISRC did not form a single cohesive population.

**Figure 6.**
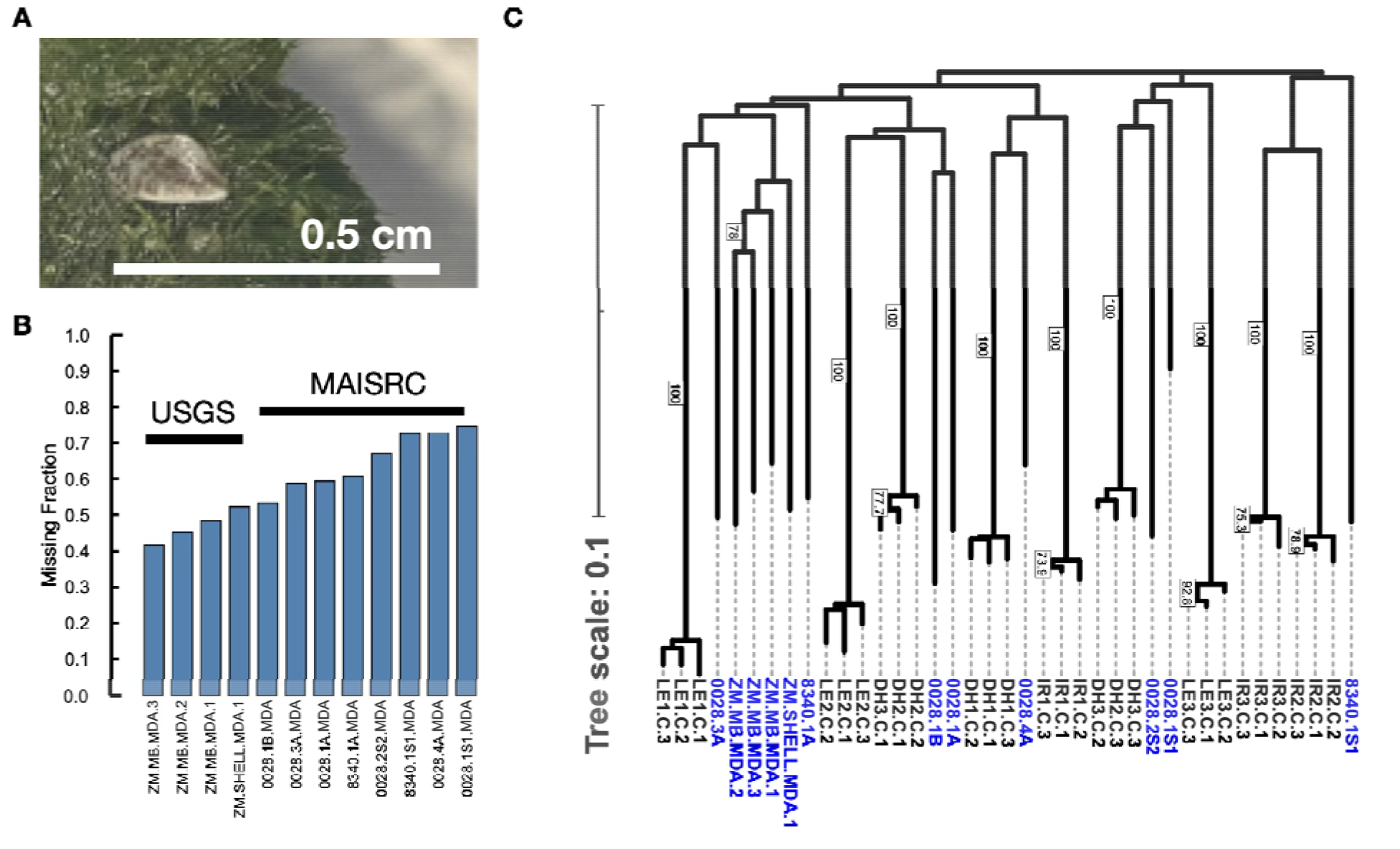
Genotyping Data from Zebra Mussels Found in Aquarium Moss Balls. (A) Photograph of a zebra mussel confiscated from aquarium moss balls. (B) Missing genotyping fraction for moss ball samples against zebra mussel reference genome. C) BioNJ phylogenetic tree generated from genetic bitwise distances. Labels at nodes indicate bootstrap support values based on 1,000 replicates. This dendrogram compares moss ball samples (sourced from MAISRC and USGS) against established reference populations (DH, IR, and LE) using 1,207 markers across 248 loci (MDA samples in blue, outgroups in grey).

## Discussion

Currently, aquatic ecosystems are experiencing dramatic changes due to the introduction of aquatic invasive species (AIS), which is largely driven by human activity. The impacts of AIS are reflected in the disruption of the ecological goods and services that water resources provide. Specifically, the zebra mussel has been identified as highly detrimental to freshwater ecosystems in North America, with a massive acceleration of infestations occurring between 2003 and the present, leading to substantial damage to native species. In a previous study employing high-resolution population genomics, we mapped the genetic signatures of zebra mussel populations across numerous inland lakes in Minnesota and compared them against potential source waters, such as the Great Lakes and the Upper Mississippi River basin. This established Lake Superior, the Upper Mississippi River, Lake Erie, and several inland lakes as the top-ranked inferred source populations for Minnesota inland lakes between 2003 and 2018^22^. This study shed light on the potential mechanisms underlying geographic expansion and provided genomic surveillance data to manage the continuing spread of zebra mussels. However, applying such an approach has been limited to well-established infestations where adult mussels can be readily obtained. In many cases, zebra mussels are first detected in a newly infested water body through the identification of veliger larvae in plankton tows. Thus, methods for genotyping low-biomass veliger samples are needed in order to provide rapid information for early-stage infestations, where timely decisions are critical for investigating sources and preventing spread.

Here, we implemented an MDA protocol to amplify genomic material from both low-concentration adult and single-larva zebra mussel DNA. Overall, we observed that MDA-amplified DNA can be used for a range of molecular applications such as PCR, GBS, or WGS. For WGS, we observed that MDA did not affect quality metrics such as Q-scores, the percentage of aligned reads, coverage, or the total variant counts for both zebra mussel and Drosophila samples (Figure 2).

Previously, we have observed well-differentiated zebra mussel populations in the Great Lakes and Midwestern United States, exemplified by the DH, IR, and LE populations. We generated genotyping data from diluted and MDA-treated DNA from these three populations (Figure 3). Sequencing data resulted in consistent quality metrics, including mean quality, GC content, deduplication percentage, and alignment rates, between treated and control samples. For the genotyping data, a lower fraction of missing genotypes (∼25%) and a smaller number of total variants (∼5000 fewer variants) were observed for MDA-treated samples, but the total number of called SNPs remained consistent. Given that GBS analyses primarily rely on the detection of SNPs, the stable count of this variant type underscores the feasibility of using this approach for SNP-based genotyping. To determine the impact of MDA on population structure, we calculated genetic distances using the BioNJ method for these three zebra mussel populations (Figure 3D). We observed that control samples formed distinct, tight clusters alongside their corresponding MDA-treated counterparts, reflecting reproducibility across both technical and biological replicates. In addition, the control and MDA-treated samples grouped reliably with their correct source populations, demonstrating consistency to the untreated controls.

When executing whole-genome amplification on single-cell or ultra-low-input DNA, MDA has been associated with several technical limitations. Specifically, potential sequence-dependent GC bias, uneven coverage distribution (locus dropouts), and compromised variant accuracy driven by allelic dropout or chimera formation^56–58^. Here, we observed a 0.031% GC dropout for WGS of MDA-treated adult DNA from the Minnetonka population (Figure 2J), which indicates a low percentage of misaligned reads linked to regions with high GC content. While low-input MDA typically disrupts coverage uniformity, pairing MDA with a reduced-representation method like GBS appears to buffer against stochastic whole-genome fluctuations.

Regarding variant accuracy, while the single-cell genomics literature frequently reports distorted heterozygosity due to preferential allele amplification^59,60^. In the case of GBS, distortions in allele frequency are probably more likely to result from PCR biases in library amplification than from the MDA process. Here, our approach yielded highly stable SNP counts for both WGS (Figure 2F) and GBS data (Figure 3C). The compact clustering between control and treated samples (Figure 3D) confirms that any minor shifts in total variant counts (Figure 3C) did not introduce false genetic distances or compromise the population genomic signal, suggesting its potential for high-fidelity, low-biomass surveillance.

After establishing optimal conditions for MDA, and assessing its effect on sequencing and genotyping data generation from diluted adult zebra mussel DNA, we moved forward with genotyping single veliger samples from Lake Minnetonka (Figure 4A). We have determined that DNA extraction from a single veliger can yield in the order of 1 ng, an amount that is within the input mass functional range for MDA (Figure 2C). As observed for diluted adult DNA, sequencing metrics for veliger DNA were comparable with adult controls (Figure 4B-4C). When comparing the genomic distances between the three control populations (Duluth Harbor, Illinois River, and Lake Erie) and Lake Minnetonka, most of the Minnetonka adults and veligers grouped together within a single cluster, and this population was differentiated from DH, IR, and LE (Figure 4D).

In the genetic trio experiment, we confirmed that offspring genotypes followed expected parental transmission (Figure 5G), consistent with Mendelian inheritance, further confirming the accuracy of MDA-derived genotyping data. We did not compute genome-wide Mendelian error rates because variable coverage constrained the number of trio-informative loci; thus, this inheritance analysis is primarily illustrative. In addition to population genomic studies, the development of low biomass genotyping technologies have the potential to facilitate the analysis of engineered genetic modifications or payloads (e.g., delivered via CRISPR-Cas9 injections). Because early-stage zebra mussel veligers are microscopic and yield very small quantities of DNA, verifying offspring genotypes would otherwise require rearing them for months to later developmental stages, a notoriously difficult and time-consuming process in a laboratory setting. By demonstrating that MDA can sufficiently amplify veliger DNA to track Mendelian inheritance patterns directly from parents to 5-day-old offspring (Figure 5G), this study lays the groundwork for a rapid screening pipeline. Thus, MDA can potentially be employed to confirm the transmission of modified alleles and accelerate the validation workflow for novel genetic biocontrol strategies.

The generation of genotypic data from confiscated aquarium moss ball zebra mussel samples demonstrates the utility of MDA for GBS-based ecological forensics, even when sample preservation is suboptimal. Preserving tissues in frozen water, with the accompanying freeze-thaw cycles, can lead to cell lysis, DNA degradation, and cross contamination, which may explain the observed low alignment rates and high fraction of missing genotypes in these samples. However, the recovery of 1,207 markers across 248 loci indicates that GBS can still capture genomic data to perform basic population structure analyses. The genomic divergence of the USGS samples is significant from an invasion biology perspective. Because these samples formed a separate clade and did not cluster with the reference populations (DH, IR, and LE), it is possible that this unique genetic signature could be used to identify future infestations resulting from an introduction related to these imported aquarium moss ball samples. Future studies tracking aquarium trade interceptions should prioritize immediate tissue preservation or DNA extraction to minimize technical artifacts.

Here, we demonstrated that pairing MDA with GBS has the potential to overcome the biomass limitations of genomic surveillance for early-stage zebra mussel infestations. By amplifying low-input DNA, including samples from single veligers at a ≤1 ng threshold, the MDA protocol yielded reproducible genomic variant data suitable for population assignment. We acknowledge that orthogonal validation using non-restriction-based genotyping technologies, such as targeted amplicon sequencing or microsatellite panels, will be required to establish locus-level genotype accuracy, particularly for applications involving degraded samples or forensic assignment. Nevertheless, the ability to generate genotyping information from veliger larvae changes the logistics of field sampling and zebra mussel genomic surveillance, particularly for early detection.

In conclusion, this study uses zebra mussels to benchmark a robust workflow for genotyping of low-biomass samples. We anticipate that this approach could be readily adapted to diverse organisms and applications, including population genomics of insects, microbes, or early life stages, non-destructive sampling of live organisms, and minimally invasive sampling from precious samples, such as museum specimens.

## Acknowledgments

The authors thank Krista Espelien, Ben Minerich, and John Gerritsen for the collection of zebra mussels and plankton tows. We also acknowledge Nick Phelps from the MAISRC, and Stephen Spear from the U.S. Geological Survey Upper Midwest Environmental Sciences Center, for providing the moss ball samples. We thank Kenny Beckman and Michelle Riehle for helpful discussions. We acknowledge Noah Strom from the University of Minnesota Genomics Center for sharing code for the GC Bias calculations and comparisons. We thank our colleagues in the University of Minnesota Genomics Center (RRID:SCR_012413), as well as the entire UMGC Innovation Lab staff. The authors acknowledge the Minnesota Supercomputing Institute (MSI) at the University of Minnesota for providing resources that contributed to the research results reported within this paper. URL: http://www.msi.umn.edu. Stocks obtained from the Bloomington Drosophila Stock Center (NIH P40OD018537) were used in this study. Funding was provided by a grant from the US Geological Survey (FY24 WRRA AIS Grant #G25AP00161-00).

## Author Contributions

Victor H. Hernandez Elizarraga: Conceptualization; Investigation; Data curation; Visualization; Writing – original draft. Lindsey G. O’Brien: Writing – reviewing & editing; Investigation. Michael A. McCartney: Investigation; Writing – reviewing & editing. Daryl M. Gohl: Conceptualization; Writing – reviewing & editing; Supervision; Project Administration

## Data Availability Statement

The raw sequencing datasets generated and analyzed during this study have been deposited in the National Center for Biotechnology Information (NCBI) Sequence Read Archive (SRA) under BioProject accession number PRJNA1519718.

## Supplementary Figures

**Supplementary Figure 1.**
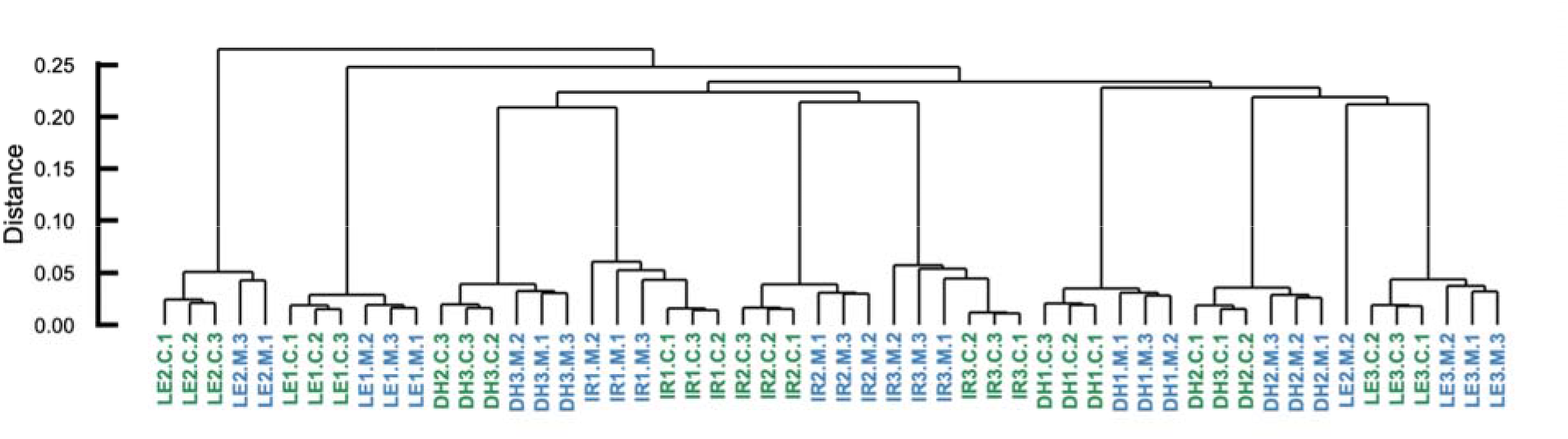
Genomic distance analysis using identity-by-state genetic distances. Dendrogram of zebra mussel adult DNA from control (C, green) and MDA-treated (M, blue) replicates from the same biological source.

**Supplementary Figure 2.**
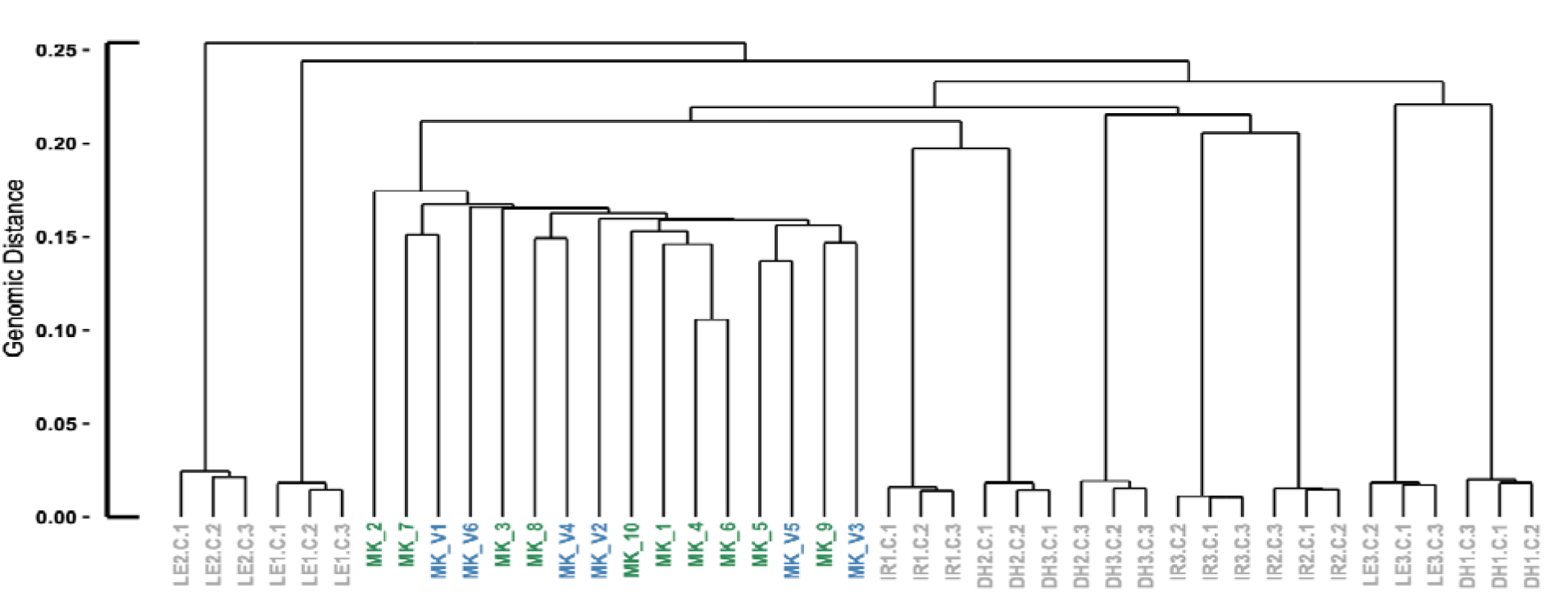
Calculated identity-by-state genetic distances for Minnetonka (MK) adults and veligers (MK_V) alongside the three reference populations from Figure 2.

**Supplementary Figure 3.**
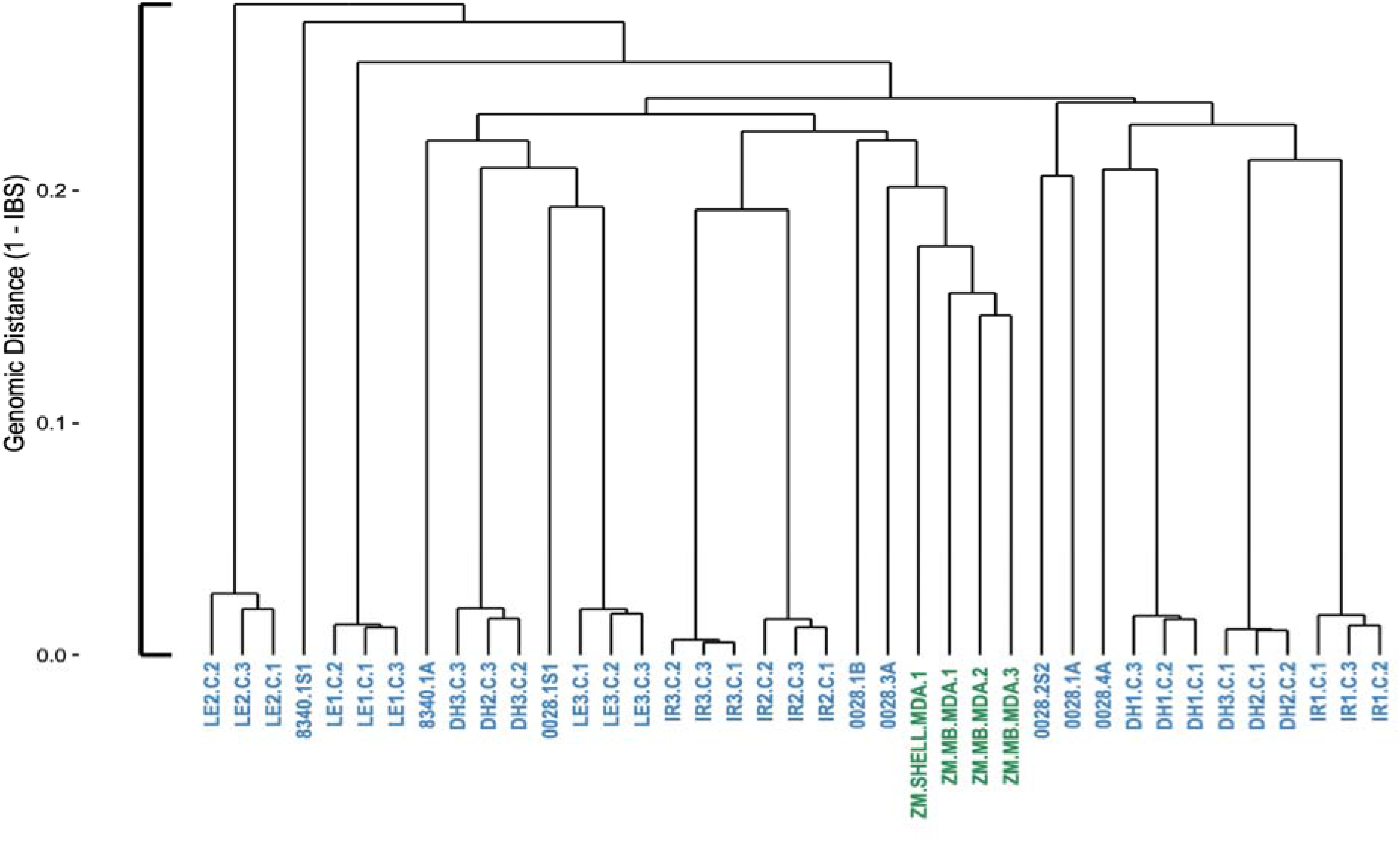
Genomic distance analysis, using identity-by-state, comparing moss ball samples (sourced from MAISRC and USGS) against established reference populations (DH, IR, and LE) using 1,207 markers across 248 loci (MDA samples from USGS in green).

**Supplementary Figure 4.**
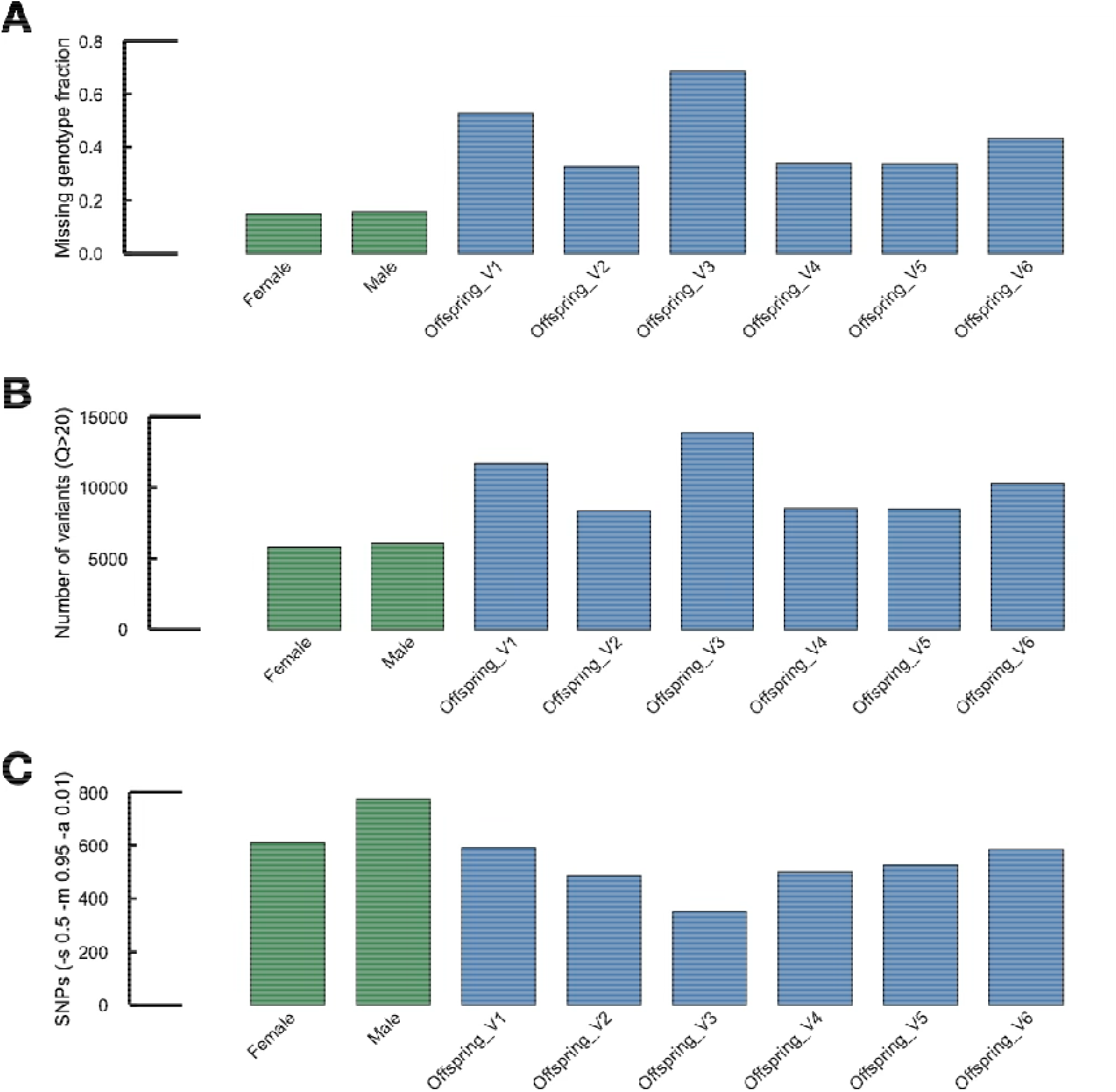
MDA sequencing performance on zebra mussel offspring. (A) Missing genotyping fraction. (B) Total number of variants. (C) Number of SNPs. Green indicates parents, and blue indicates offspring treated with MDA.

